# Efficient decision-makers evaluate relative reward per effort

**DOI:** 10.1101/2022.05.31.494175

**Authors:** Jan Kubanek

**Affiliations:** University of Utah, 36 S Wasatch Dr, Salt Lake City, UT 84112

## Abstract

Understanding how humans and animals can make effective decisions would have a profound impact on economics, psychology, ecology, and related fields. Neoclassical economics provides a formalism for optimal decisions, but the apparatus requires a large number of evaluations of the decision options as well as representations and computations that are biologically implausible. This article shows that natural constraints distill the economic optimization into an efficient and biologically plausible decision strategy. In this strategy, decision-makers evaluate the relative reward across their options and allocate their effort proportionally, thus equalizing the reward per effort across their options. Using a combination of analytical and simulation approaches, the article shows that this strategy is efficient, providing optimal or near-optimal gain following a single evaluation of decision options. The strategy is also rational; satisficing and indifferent decision-makers are found to perform relatively poorly. Moreover, the relativistic value functions underlying this efficient strategy provide an account of time discounting, effort discounting, and resource discounting in general.

---

Neoclassical economics provides a powerful apparatus that dictates how humans and animals should allocate their time, effort, or monetary resources^1, 2^. A hallmark of this normative set of theories is that decision-makers maximize a criterion—individuals allocate their resources to maximize their total utility or payoff^3–5^.

Albeit normative, this set of theories has faced major criticisms^6–8^. The most severe is that there is no prescription of the strategy that decision-makers should take to maximize their total utility^9^. To attain a maximum, it is assumed that decision-makers can evaluate decision option in a near-limitless manner and perform calculus in multidimensional space^10–12^. Moreover, the apparatus requires an absolute definition of utility^13, 14^.

This article addresses these limitations by constraining the economic framework with natural considerations. By doing so, it distill the economic framework into a simple yet powerful strategy that provides efficient and rational decisions.

## Results

The normative economic framework is found to be simplified using two specific natural considerations (Methods). These considerations include 1) the law of diminishing marginal utility and 2) casting utility and effort in relative terms. The incorporation of these considerations into the economic framework provides a simple yet powerful decision strategy: a proportional allocation of relative effort to relative utility of each decision option, equivalent to evaluating relative reward per effort (Methods). This formalism is related to the matching law^39–42^, but unlike the matching law, which only describes behavioral equilibrium, this effort allocation rule constitutes a decision strategy taken by decision-makers at each decision time. Given the structural relationship to the matching law and its instant deployment, the strategy is referred to as “Instant Matching” (IM). The Methods section shows analytically that a single evaluation of decision options using IM provides an optimal or near-optimal gain. Given this finding, decision-makers that use this strategy are referred to as “efficient decision-makers”.

The analytical findings of IM efficiency are herein validated through simulations that probe a large space of choice situations. Specifically, the simulations include 2, 3, and up to 10 options, each with distinct, randomly generated relationship between relative utility and relative effort, for a total of 9000 choice situations (see Methods).

### Efficiency of Instant Matching

The efficiency of IM is evaluated by comparing the total gain of decision-makers using IM (“efficient decision-makers”) with the maximal possible gain of theoretical economic agents (see Methods for details). **Fig. 1** shows the result. Efficient decision-makers obtained an average 90.2% of the total possible gain across the choice situations tested. Crucially, they collected this level of gain following a single evaluation of their options. In contrast, the theoretical economic agents, whose computations involve multidimensional optimization, require an average of at least 16 evaluations of the decision options to reach the same level of performance (**Fig. 1**, gray).

**Figure 1.**
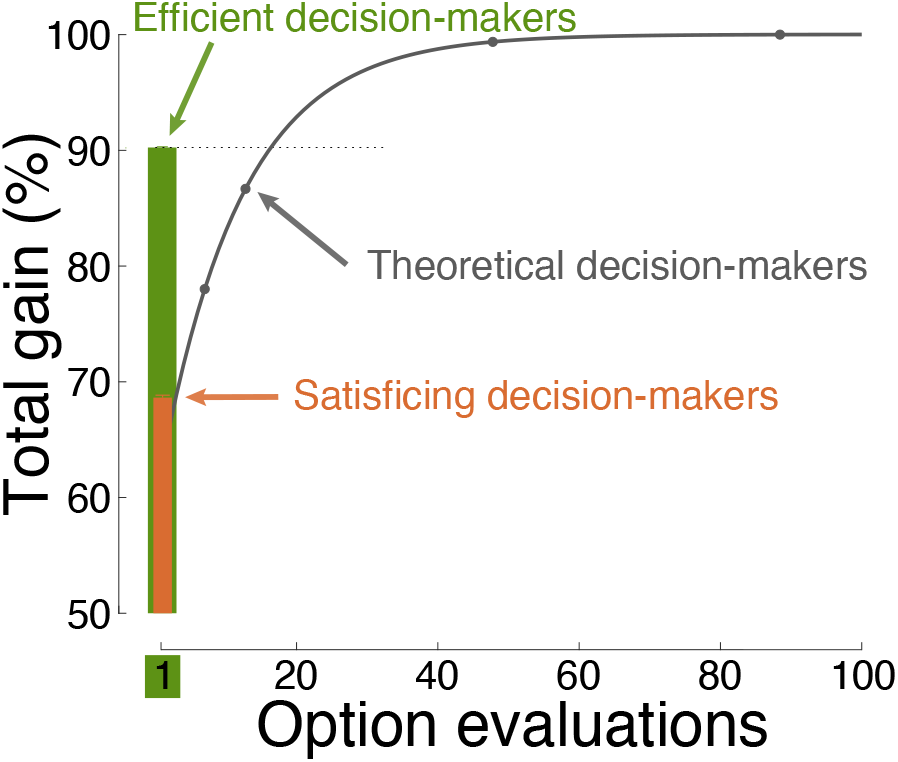
Efficient decision making following a single evaluation of decision options. The figure compares the performance of three kinds of decision-makers. The first kind (gray) performs multidimensional optimization over the decision options (see Methods). The second kind (green) uses Instant Matching. The third kind (orange) uses a satisficing strategy, putting all resources into the most desirable option. The decision-makers faced a total of 9000 simulated choice situations, involving up to 10 options, each with distinct contingencies between utility and effort (Methods). The maximum gain (the value of 100%) was established by allowing the multidimensional optimization algorithm to converge. Instant Matching (green) provides a high gain while requiring a single evaluation step. Here and henceforth, plots show mean±s.e.m. values. Here and elsewhere, the error bars are often smaller than the symbols indicating the mean.

To put this result into natural perspective, consider a foraging animal that must decide how to allocate her presence across several patches of food. If she was (and could) use the economic framework (Eq. 1), she would have to visit and forage in each patch at least 16 times to make a decision of quality comparable to IM. IM allows biological decision-makers to make effective decisions in a single step, i.e., efficient decisions.

### Rationality of Instant Matching

The rationality of IM is evaluated with respect to the performance of satisficing and indifferent decision-makers. A satisficing decision-maker does not perform the reward per effort valuation of Eq. 5, and instead puts all effort or resources into the option that provides the highest reward. **Fig. 1** (orange) shows that this approach degrades performance. On average, satisficing decision-makers only harvested an average of 68.6% of the total possible gain across their options. The difference between the rational and the satisficing decision-makers is substantial and highly significant (*t*_8999_ = 103.84, *p* = 0, two-sided t-test).

In IM, relative effort is proportional to relative reward, i.e., 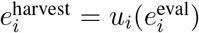, which can also be stated as 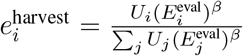, where *β* = 1. Varying the exponent enables us to test the performance of distinct kinds of decision-makers. For *β >* 1.0, a decision-maker’s behavior is drawn to rewarding options, producing satisficing behavior in the limit^13^. On the other hand, a decision-maker with *β <* 1.0 is relatively indifferent to reward. **Fig. 2a** shows that *β* = 1.0 indeed lies near the optimum, with *β* = 1.0 and *β* = 1.2 producing a 90.2% and 90.5% gain, respectively. In comparison, the satificing and indifferent strategies, albeit equally rapid, are relatively ineffective.

**Figure 2.**
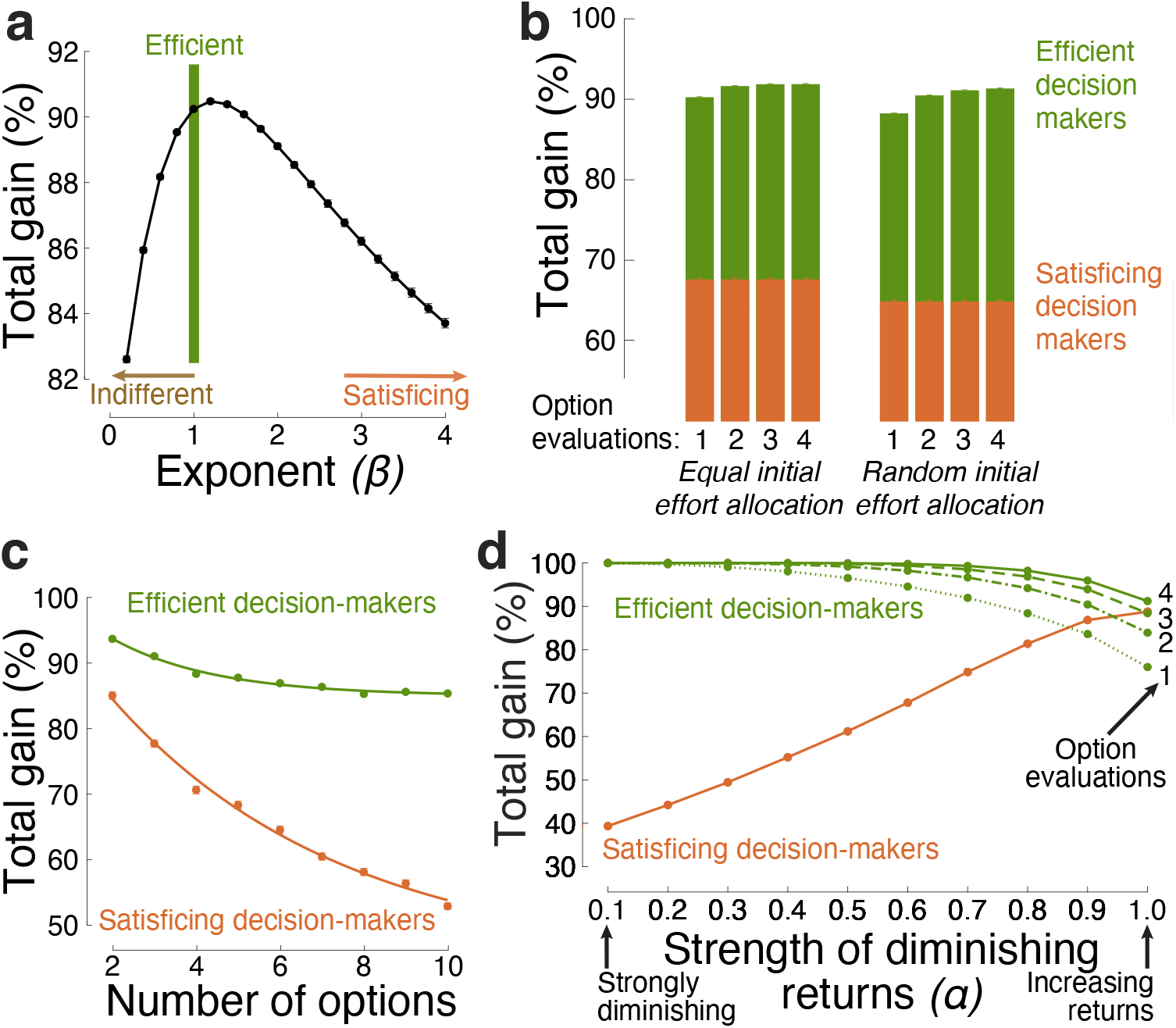
The performance and robustness of Instant Matching. **a**, Gain as a function of the level of the exponent *β* in 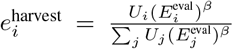. IM uses *β* = 1, i.e, relative effort is allocated in proportion to relative reward. Decision-makers who are indifferent to reward (*β* ≪ 1) and decision-makers who are strongly attracted to reward (*β* ≫ 1) suffer a performance loss. Here and elsewhere in this figure, the same 9000 decision situations as in **Fig. 1** were used. **b**, Decision-makers must evaluate decision options at least once to estimate their initial worth. In this initial evaluation, decision-makers may distribute their effort across the options equally (left bars). However, such distribution, even though desirable (see Methods) may not always be feasible. The right bars show that distributing initial effort across options entirely randomly provides a comparable level of performance for IM. This more realistic, random initial distribution is used henceforth (**c** and **d**). In both cases, the options can be evaluated iteratively, i.e., 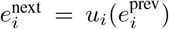, to increase performance further. **c**, Same data as in **Fig. 1**, separately for a specific number of decision options. The curves represent exponential fits. **d**, Performance as a function of the level of diminishing marginal utility. To evaluate the full spectrum from strongly diminishing to not at all diminishing (linearly increasing), this analysis specifically used the power contingency *U* (*E*)= *U*_*m*_*E*^*α*^. The abscissa shows performance as a function of 10 evenly spaced values of *α* (see Methods). Like in **(*b***, the decision strategy 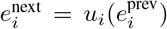 can be applied iteratively. Performance curves for up to 4 iterations are shown in distinct line styles. The satisficing strategy (orange), by definition, does not improve with additional iterations.

IM can be applied in more than one evaluation to increase the total gain further. To do so, the allocation 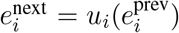 is repeated a desired number of times. **Fig. 2b**, left green bars, shows that a second evaluation increases the average performance from 90.2% to 92.0%. The performance saturates in about the third and fourth evaluation, reaching 92.3% of the possible maximum. In comparison, the performance of the satisficing strategy does not, by definition, improve with additional evaluations (**Fig. 2b**, left orange bars; see also **Fig. 3b** for a specific example).

**Figure 3.**
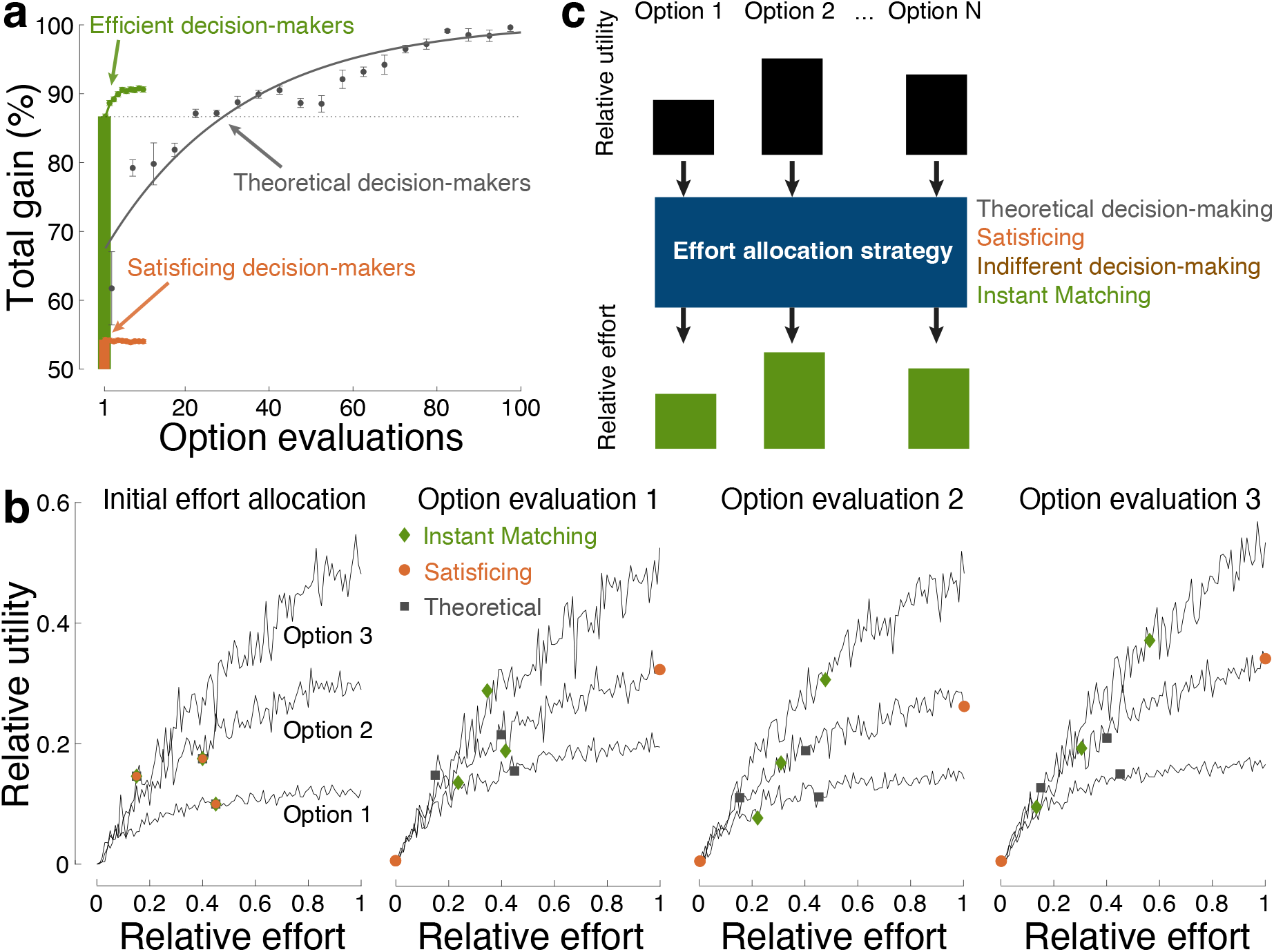
Efficient decision making in stochastic environments and graphical summary. **a**, Same format as in **Fig. 1** for decision situations in which each option provided a distinct rate of reward, according to variable interval schedules (Methods). Performance is shown over 10 (250) consecutive evaluations of the decision options for the efficient and satisficing (theoretical) decision-makers. Each strategy initiated with a random allocation of effort across the choice options. **b**, Example of effort allocation in a three-option decision situation, as a function of the individual evaluations. The steps of the individual strategies are shown as colored symbols (see inset). All strategies started with the same random initial allocation of effort. The VI schedule provides distinct sets of binary rewards on each evaluation; hence the distinct relationships between utility and effort on each evaluation. **c**, Summary. This study investigated how decision-makers can efficiently allocate relative effort to relative utilities of their choice options. The study specifically focused on the decision strategies (blue box); four major strategies were evaluated across a large space of decision situations. Instant Matching uniquely provides rapid convergence to high performance and provides an explanation for time and effort discounting (**Fig. 4**).

Thus far, we have assumed that decision-makers allocate effort or expenditures equally when sampling the initial worth of their options (Methods). However, equal sampling may be impractical or impossible in many choice situations. The following analysis reveals that IM is robust with respect to a particular initial distribution 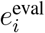. Specifically, **Fig. 2b**, right green bars, shows that decision-makers who distribute their initial effort 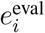 entirely randomly (e.g., **Fig. 3b**) can still apply the same strategy with comparable level of efficiency. In addition, the performance increases even more steeply by readjusting effort in response to additional option evaluations 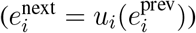. On the fourth evaluation, the performance reaches 91.6%. A two-way ANOVA with factors evaluation number (*I*) and initial effort distribution (*D*) detected a significant effect of *I* (*F* (3, 71992) = 672.42, *p* = 0), a significant effect of *D* (*F* (1, 71992) = 715.82, *p* = 6.4 × 10^−157^), and a significant interaction between the factors (*F* (3, 71992) = 51.38, *p* = 3.6 × 10^−33^). The satisficing strategy does not depend on the number of evaluations (**Fig. 2b**, orange) but is sensitive to the initial allocation of effort (*F* (1, 71992) = 300.10, *p* = 4.3 × 10^−67^).

IM is robust also with respect to the number of decision options. **Fig. 2c** shows that decisionmakers using IM perform well even when faced with a relatively high number of options. IM attains an average 93.7% gain for all situations that involved 2 options. The strategy has an exponential decay of 2.6% per option, and achieves a 85.3% gain for 10-option decisions. In contrast, satisficing decisionmakers fall short when facing an appreciable number of options. This is expected because putting all effort into the seemingly most rewarding option leads to an opportunity loss (e.g., **Fig. 3b**). The satisficing strategy shows an exponential decay of 3.9% per option, and attains a mere 52.9% average gain in 10-option situations.

The law of diminishing marginal utility, which was harnessed to derive IM, may not hold in all situations. In certain situations, a payoff may increase indefinitely with the invested effort. To investigate the effect of this possibility on IM, let us parameterize the strength of diminishing returns using the power function, *U* (*E*)= *U*_*m*_*E*^*α*^, where *α* is the tested parameter. For *α* = 0.1, the function exhibits strongly diminishing returns (**Fig. 5a**). On the other hand, for *α* = 1.0, gain increases linearly with effort and never saturates. **Fig. 2d** shows the dependence of the total gain on this parameter. As expected, IM, based on the law of diminishing marginal utility, shows high performance when the choice options obey the law. The satisificing strategy is particularly inadequate in this case. Satisficing is blind to the law of diminishing marginal utility: A satisficing decision-maker puts all effort or resources into an option whose payoff eventually saturates, thus presenting an opportunity loss (e.g., **Fig. 3b**). As expected, the performance of IM / the satisficing strategy is lower / higher in choice situations in which the law of diminishing marginal utility does not hold (*α* = 1.0). Nonetheless, IM can outperform the satisficing strategy in such cases, when applied iteratively. In particular, following the fourth evaluation, the efficient strategy attains a 91.2% gain, compared to a 88.8% gain of satisficing. The same principal observation is made for functions other than the power function (**Fig. 5b**).

Thus far, the performance of IM and its counterparts was evaluated in deterministic decision environments (**Fig. 5a**). To evaluate their performance in stochastic decision situations, the strategies were tasked to make choices in environments in which individual rewards were scheduled randomly, according to the variable interval schedules (see Methods). The decision-makers integrated the individual binary rewards over a defined period of time using reinforcement learning (see Methods), and allocated effort according to the estimated utility of each option. The rate of reward occurrence for each option was again randomized across 1000 distinct decision situations, each involving 2, 3, and up to 10 decision options.

IM was found to be efficient also in these stochastic situations (**Fig. 3a**). Following a single evaluation of the decision options, IM harvested an average of 86.7% of the total possible gain. In comparison, the theoretical decision-makers required an average of 29 evaluations of the decision options to reach the same level of performance. Satisficing produced an average 54.1% gain in the stochastic environments. As in the deterministic cases assessed previously, the performance of IM improved with repeated application of its 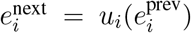 strategy, reaching an average 90.0% gain on the fourth evaluation.

**Fig. 3b** illustrates how IM rapidly reaches high gain under limited representational and computational resources. Following an initial assessment of the relative worth of the available options (left panel), IM (green diamonds) allocates effort in proportion to the relative worth, i.e., 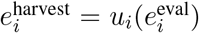. The second panel shows that this step readily results in harvesting the highest reward from the richest option (Option 3), as desired, while also gaining substantial reward from the relatively less valuable options. Repeated application of this simple strategy (third and fourth panels) leads to additional gain. In comparison, the satisficing strategy (orange circles) settles with the seemingly richest option, and can miss the truly richest option as shown here. Even if equal initial effort allocation allowed this strategy to identify the richest option, the diminishing returns of the choice environment would present an opportunity loss with respect to other options. The theoretical strategy (gray squares) samples the decision options based on gradients computed within the mutidimensional space of decision options. This approach requires a large number of option evaluations to reach the performance level of IM, as apparent in **Fig. 3a**.

## Discussion

This study derives an efficient decision strategy from the normative economic framework. The key finding is that efficient decisions are made by evaluating the relative reward per relative effort associated with the decision options (IM strategy). IM is efficient in that it provides optimal or near-optimal gain following a single evaluation of the decision options. IM is also rational in that its satisficing and indifferent counterparts perform relatively poorly.

IM is simple yet highly efficient. In particular, IM was benchmarked across 9000 decision situations of up to 10 decision options, and harvested around 90% of the maximum possible gain following a single evaluation of the decision options. Moreover, in human settings, in which the worth of the options may be known or can be estimated (e.g., price), IM allows individuals to allocate effort (e.g., money) readily, without any prior sampling. Thus, IM is efficient, combining high performance with rapid and simple evaluation.

IM has been considered by ecologists as a candidate for an evolutionarily stable strategy^15, 16^. Referred to as the *relative payoff sum* in behavioral ecology, a population using this strategy is believed to be uninvadable by a mutant with a different strategy^15, 17^. A proof was attempted^15^ but remains elusive^18^. The present study shows that this strategy stems from the maximization framework of neoclassical economics, and shows an analytical relationship of IM to this normative framework. Critically, analytical derivations and simulations show that the strategy is efficient, providing optimal or near-optimal gain in a single evaluation of the choice options in situations in which the law of diminishing marginal utility holds (**Fig. 2, Fig. 5**). Given that foraging efficiency is a critical optimization variable in ecology^19^, this study supports the notion of this strategy as evolutionarily stable, under the assumption that the law of diminishing marginal utility is a prevalent natural phenomenon.

As in ecological models^15, 17^, this study specifically focuses on the decision strategies used to allocate effort to available options (blue box in **Fig. 3c**). How decision-makers can represent the relative worth or relative utility of decision options has been addressed in many previous studies in ecology, computer science, and neuroscience^20–23^. Notably, the relativistic nature of *u*_*i*_ and *e*_*i*_ enables these variables to be subjective. So long as the underlying representations are comparable by the brain (e.g., through relative neuronal firing rates), the *e*_*i*_ = *u*_*i*_ strategy provides an efficient allocation of the decision-maker’s resources. Moreover, *u*_*i*_ can incorporate additional factors such as the probability of an outcome, as in prospect theory^24^. A previous study^25^ demonstrates that prospect theory’s incorporation of probabilities into utilities does not change the relationship between the differential formulation of Eq. 1 and the fractional formulation of Eq. 2, which is crucial for IM.

The value representations underlying IM align with the empirically established value functions in psychology, including time and effort discounting. This study provides a teleological reason for such representations—efficiency with respect to limited resources, including time and effort, and their saturating returns. As a consequence, efficient decision-makers evaluate the reward rate (i.e., reward per time), the reward per effort, or, most generally, the reward per the applicable resource across their options. An example of the reward rate valuation are decisions concerning job choice based on salary (i.e., monetary income per unit of time).

Decision-makers are known to weigh benefits against costs, or rewards against effort. Yet, how effort should be incorporated in current models of decision-making in economics^26^, ecology^27^, and psychology^28, 29^ has been unclear. This study shows that individuals can make efficient cost-benefit decisions by evaluating the relative reward *u* per relative effort *e* of each option, i.e.,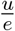. This formulation increases with the benefit of an option (*u*), decreases with the cost of an option (*e*), and the relativistic expression does not require any free parameter.

The derived equations provide a normative insight into the parameter *k* used in studies of time and effort discounting. Specifically, 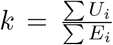 serves to convert absolute utilities and efforts into relative quantities. Since *k* represents the total possible gain per total available resource, it provides specific predictions regarding the rates of discounting for each individual and each decision domain. For example, consider a decision-maker who can devote a limited amount of time to a task. The lower the amount of available time, the higher the *k*, and so the more steeply the individual discounts a gain with respect to a time investment 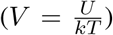. Reversely, if a resource such as time is abundant, a payoff is discounted relatively little. Thus, the discounting factor incorporates the scarcity or the abundance of each decisionmaker’s resources.

The alignment of the value functions associated with IM with value functions empirically obtained in psychology provides further validation of the considerations used to arrive at IM (**Fig. 4**). First, decision-makers have limited resources in the form of time and physical and mental energy. Second, expenditures of these resources commonly reach saturation or diminishing returns^30–33^. And third, decisions involve relative rather than absolute representations.

**Figure 4.**
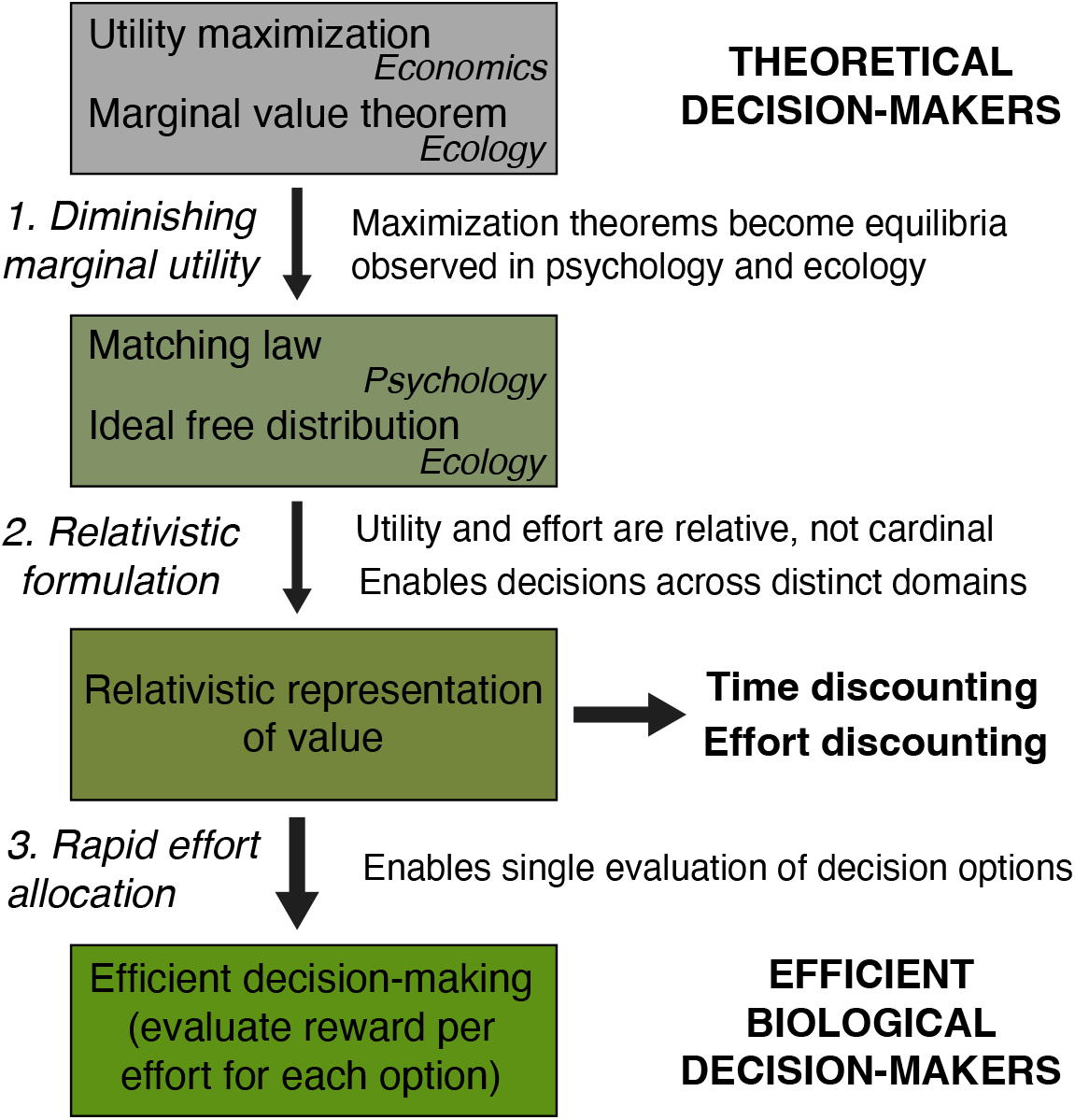
Framework for efficient biological decision-making. Steps, detailed formally below, that relate the maximization framework of economics (gray, top box) to value representations in psychology and an efficient biological decision framework (green, bottom box). These steps take into account natural and biological phenomena and constraints (see text for details).

**Figure 5.**
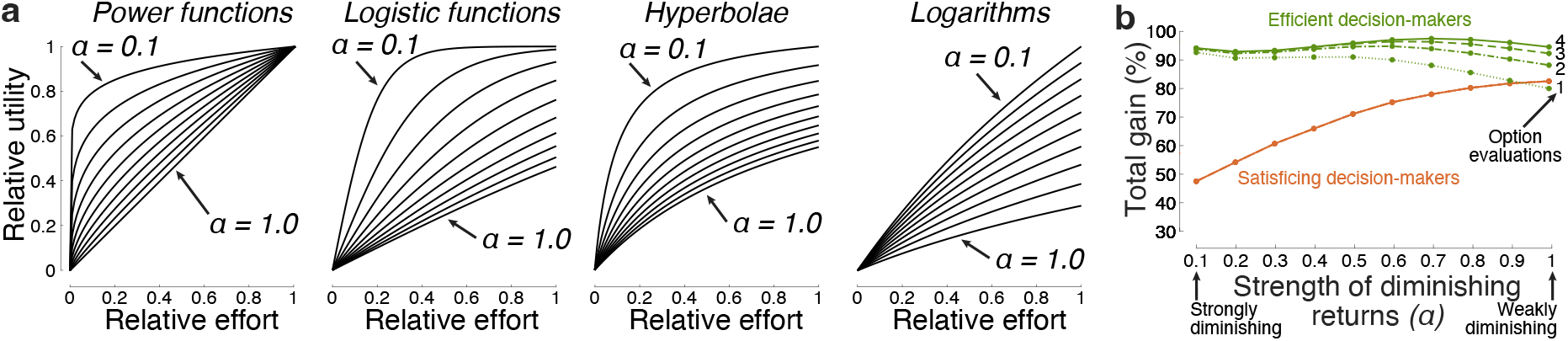
Performance as a function of the level of diminishing marginal utility. **a**, The utility-effort contingencies tested. Power, logistic, hyperbolic, and logarithmic functions in the forms 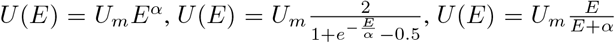, and 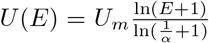, respectively, were used. In these functions, the variable *α* quantifies the level of diminishing marginal utility, and ranged from 0.1 (strongly diminishing) to 1.0 (least-diminishing; linear for the power function), in steps of 0.1. The individual plots are scaled by these 10 values of *α* in the indicated order. In addition, the variable *U*_*m*_ in these functions scales the magnitude of each contingency, ranging from 0 to 1. **b**, Average performance of Instant Matching and its satisficing counterpart as a function of the level of diminishing marginal utility. Same format as in **Fig. 2d**, but now for any combination of the functions shown in **a**. Specifically, each option (2 up to 10; 1000 evaluations each) was randomly assigned any of the four functions with the diminishing marginal utility level *α* indicated on the abscissa. Strong diminishing returns provide an almost immediate convergence onto the optimum. Iterative application of IM greatly improves performance in cases with weakly diminishing or increasing returns (see number of option evaluations in black).

With respect to the representation of utility^10–14^, this study finds that individuals can make efficient decisions by representing utilities and efforts relativistically (Eq. 5). This has important practical implications. First, relativistic representations enable neurons in the brain to encode the worth of decision options through relative firing rates or other relative means^21^. Second, relative utilities allow individuals to make decisions across distinct domains. Specifically, relative utilities or rewards can be evaluated over comparable domains (e.g., the domain of foods, the domain of activities). Like relative utilities, relative efforts can also be evaluated over comparable domains (e.g., the domain of time, the domain of mental effort). The associated value representations, 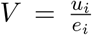, can thus be used for decisions across distinct domains. Actions are directed to options in any domain that provide high relative reward per relative effort.

IM is testable at the behavioral and neural levels. At the behavioral level, IM possesses three distinctive characteristics. First, IM obtains high reward rapidly. This characteristic can be tested in choice environments that minimize noise as an additional factor, providing performance versus evaluations plots analogous to **Fig. 1**. Second, IM allocates relative effort to relative utilities proportionally (Eq. 7). This characteristic can be tested in situations in which utilities can be measured precisely, e.g., through the volume of fluid rewards in animal experiments or money in human experiments. And third, IM allocates effort rapidly regardless of the number of options. This is because the 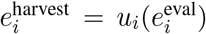 strategy is agnostic to the number of options. This characteristic can be tested by varying the number of decision options and quantifying the number of times a decision-maker evaluates the options. At the neural level, IM only requires the encoding of relative utilities of the recently sampled options. This relative code can be implemented using firing rates of the neuronal pools representing each alternative. Indeed, this representation has been found in the primate brain. Specifically, the relative value associated with IM, termed “fractional income,” captures firing rates of neurons in monkey area LIP^21^.

In IM, the relativistic representation of utility (Eq. 7) involves divisive normalization. Divisive normalization a common operation performed by neural circuits^34^. The specific form of this operation may be crucial for explaining attraction, similarity, and compromise effects observed in multi-alternative, multi-attribute decision environments^35, 36^. For instance, it has been found that a transformation of utilities by specific monotonic functions prior to divisive normalization can explain these behavioral effects parsimoniously^36^. On this front, monotonic transformations and divisive normalization are performed by several kinds of feedforward and feedback neural circuits^34, 37^. Nonetheless, how exactly individual attributes of decision options are encoded at the neural level should be investigated using large-scale neuronal recordings.

In summary, this study provides a decision strategy for how decision-makers can make efficient decisions based on the relative worth of their options. Optimal or near-optimal decisions are achieved by evaluating relative reward per relative effort across choice alternatives. Such decisions are made rapidly, using a single evaluation of the decision options. Moreover, the underlying value representations align with several observations in psychology, ecology, and neuroscience. This biologically plausible framework is expected to provide an anchor for understanding the efficiency and rationality of choice behavior in behavioral economics, psychology, ecology, and related fields.

## Methods

### Link between economic maximization and efficient biological decision-making

The economic maximization framework provides an analytical prescription for the highest possible gain given resource constraints^2^, and thus constitutes an essential starting point (**Fig. 4**, top). According to this framework, subjects maximize the total expected utility across their *n* options, 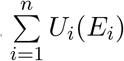. Here, *U*_*i*_ is the expected utility of an option and *E*_*i*_ the effort or expenditure required to produce that utility. The framework takes into account the decision-maker’s limited resources, such as effort or time, 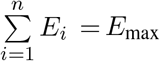. (The terms expenditure, effort, resource, and behavior are used interchangeably in this text as individuals’ means to obtain an outcome, and are all labeled *E*.) The maximization of total utility requires that effort or resources be allocated to the options such that the marginal utilities across all options are equal^25^:

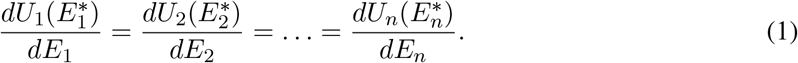

When effort embodies foraging time (*E* = *T*), an equivalent form of this equation has been presented as the *marginal value theorem* in ecology^19^.

#### Step 1. Apply the law of diminishing marginal utility

A previous study^25^ showed that the marginal fractions of Eq. 1 can be expressed as absolute fractions for an infinite number of *U* (*E*) relationships that follow the law of diminishing marginal utility^38^. This leads to the following simplification:

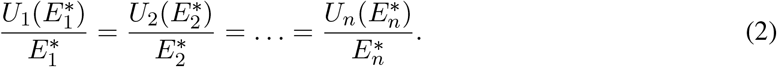

In psychology, this commonly observed equilibrium state is known as the *matching law*^39–42^. In ecology, the expenditure of a society of foraging individuals is equivalent to the number of individuals *N*_*i*_ devoted to each foraging patch. In this case, 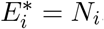, and the equation becomes the *ideal free distribution*^33, 42^,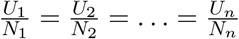.

#### Step 2. Cast utility and effort in relative terms

Albeit simpler, Eq. 2 requires decision-makers to enumerate cardinal, numeric utilities. To address this issues, observe that each of the fractions in Eq. 2 amounts to the same value. Let us denote this value as *k*. Accordingly, 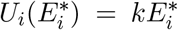, and so 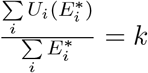. Therefore,

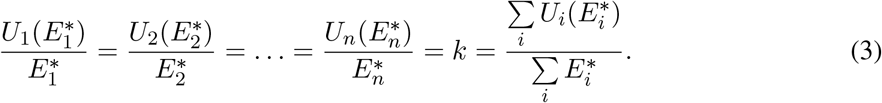

The normalization by *k* leads to a relativistic form of the matching law:

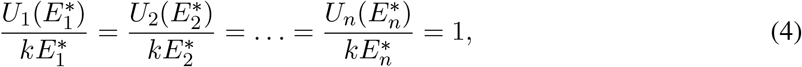

where 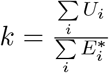 (see Eq. 3).

The normalized fractions in this equation are prime candidates for a rational representation of value. First, the expressions include both the benefits (reward) and costs (effort) associated with each option, in line with the axioms of rational choice theory^26^. Second, the values can be compared across options and are equalized in equilibrium (Eq. 4). Indeed, the value representation 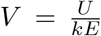 captures several phenomena in psychology (**Fig. 4**). In particular, decisions that involve time investment (*E* = *T*) and yield monetary or other outcomes *A* are characterized with 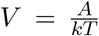. This expression embodies *time discounting*^43, 44^. Moreover, many situations require an investment of physical or mental effort *E* to obtain a reward. In these cases, the value function takes the form 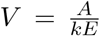. This embodies *effort discounting*^28, 29^. The Analytical derivations section proves that value functions in the form 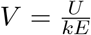 are tied to economic maximization under the law of diminishing marginal utility in a way analogous to the previously demonstrated 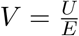 ^25^.

Critically, the normalization of Eq. 3 by 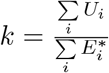 turns absolute utilities *U*_*i*_ into relative utilities 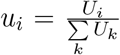. This relativistic formulation circumvents the problematic cardinal definition of utility^13, 14^, and is in this text referred to as “reward”. Furthermore, the normalization turns the absolute effort values *E*_*i*_ into relative efforts 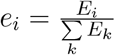. This leads to the following result:

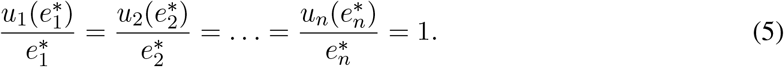

This result states that decision-makers should distribute relative efforts in proportion to relative rewards of their options:

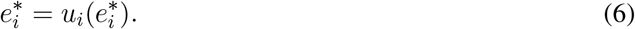

This relativistic formulation enables decision-makers to make choices across options of distinct kinds (see Discussion), and is crucial for the final step.

#### Step 3. Derive an effort allocation strategy based on the relativistic formulation

Eq. 5 embodies a relativistic form of the matching law, which represents an equilibrium state. This equilibrium is rational under the law of diminishing marginal utility^25^. This section turns the formalism into a time-by-time, local strategy that enables humans and animals to make efficient decisions. The relativistic formulation of the previous section is key to this step. In this formulation (Eq. 6), both sides of the equation, *e*_*i*_ and *u*_*i*_, are on the same relativistic scale and 2 (0, 1). This way, the ideal 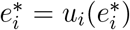 can be approximated through an iterative rule

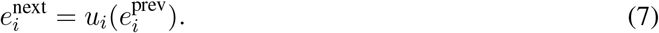

For a single evaluation of the decision options, this rule becomes

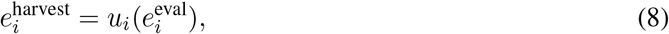

where 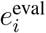 is the initial assessment of the worth of the decision options, and 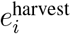 is the resulting effort allocation.

The Analytical derivations section proves that a single evaluation of the decision options provides optimal gain when the reward functions *u*_*i*_ saturate logarithmically with respect to effort. The section further shows that a single evaluation converges to an optimum for exponential *u*_*i*_ that saturate strongly with increasing effort. Finally, simulations performed in the Results section show that this strategy provides high performance also for other kinds of functions, even when assigned randomly across options, and that it can provide high performance also in situations in which the law of diminishing returns does not hold.

### Analytical derivations

#### Optimality of the relativistic formulation of the matching law

This section derives the conditions for which the relativistic form of matching, 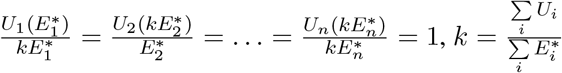, is optimal. The proof proceeds in a similar fashion as in^25^. In particular, the relativistic formulation of the matching law (Eq. 5; see also Eq. 3), which now in addition to^25^ includes *k*, is optimal when the following relationship holds for each decision option *i*:

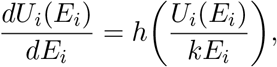

where *h* is a strictly monotonic function. For simplicity, here, let us label *U*_*i*_(*E*_*i*_) as *U* (*E*) and *E*_*i*_ as *E*.

The derivative of this equation with respect to respect to effort is

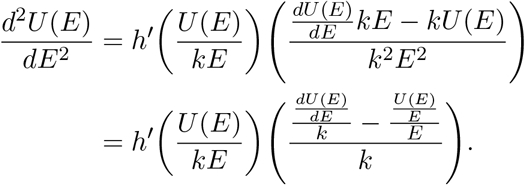

Using the above 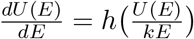, we obtain:

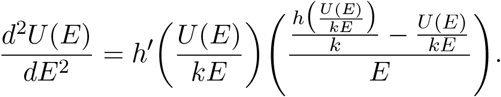

As in the main text, let the term 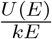 represent a value function *V*, 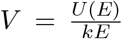. The beauty of this representation shines through the above equation. Given that *E >* 0, it is apparent that the sign of the right side of the equation is solely a function of *V* :

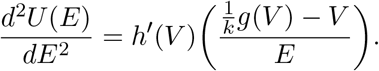

In order for a critical point to represent a reward maximum, it must be 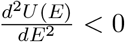 for at least one option and at least for some value of *E*^25^. To meet that requirement, and taking into account *E >* 0, there are two possibilities:

1. *h*′(*V*) > 0 and *h*(*V*) < *kV*
2. *h*′(*V*) < 0 and *h*(*V*) > *kV*

The second possibility is inadmissible because for a general *V >* 0, a function *h*(*V*) cannot be decreasing while maintaining a value above a certain line *h*(*V*) = *kV, k >* 0. This leaves us with the first set of requirements, *h*^*′*^(*V*) *>* 0 and *h*(*V*) *< kV*.

For relativistic matching, 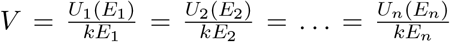 is the same for all options *i*. Thus, when 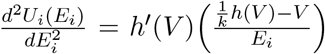 assumes a positive or a negative sign, it does so equally for all options, and the sign is governed solely by the properties of *h*(*V*). For matching yielding a reward maximum (*h*^*′*^(*V*) *>* 0 and *h*(*V*) *< kV*), it is then apparent that 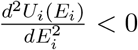 for all options and for any *E*_*i*_. Thus, all options yield diminishing returns for all values of effort.

#### Optimality of a single evaluation

This section derives a general set of functions for which Instant Matching is optimal in a single option evaluation step. Consider a general function Eq. 1 states that 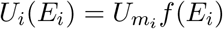. Eq. 1 states that 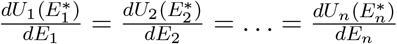. Thus, 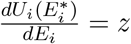, where *z* is a constant, and

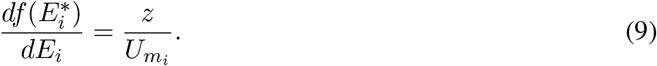

Let Instant Matching be optimal in a single evaluation step, 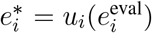, and let the initial evaluation 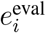 of the decision options be distributed equally. Using Eq. 4, we can also state this fact as 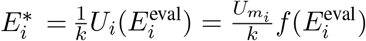. From here, 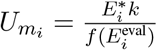. Inserting this expression into Eq. 9, we obtain

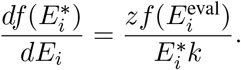

For a general function argument 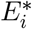, a derivative proportional to 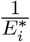 can be obtained for logarithmic functions. Thus, functions in the general form 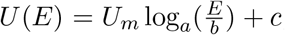, where *U*_*m*_ *>* 0, *a>* 1, *b>* 0, and *c* ∈ *R* are free parameters, provide the sought space of solutions.

Let us validate this result. When the decision-maker distributes her initial effort equally across the options, 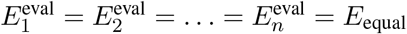, she harvests, for a general function 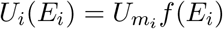.

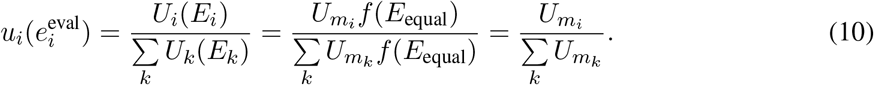

Consider now our set of solutions, logarithmic functions in the form 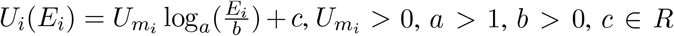. From Eq. 1, we obtain: 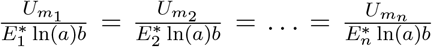. This way, 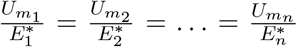. Analogously to Eq. 3, it is easy to see that 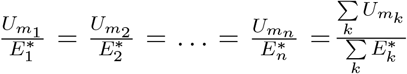. Diving all terms by the right hand side term, we obtain 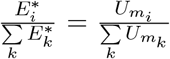, and so, relativistically written, 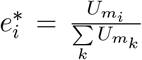. This effort allocation provides the same rewards as the initial option sampling in Eq. 10, i.e., 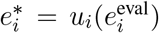. This proves that contingencies that can be characterized with a set of logarithmic functions, 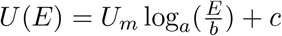, render Instant Matching optimal in a single evaluation step.

#### Efficiency and diminishing marginal utility

Let us now work toward a more general conclusion. Consider decision options for which gain is characterized by a set of power functions, 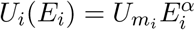. From Eq. 1, we obtain: 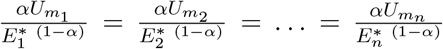. This can be rewritten as 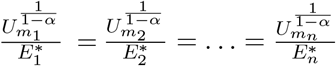. Analogously to the previous case, this equation is also equal to 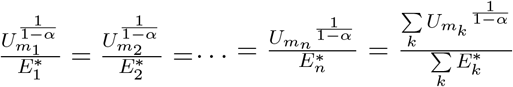. Dividing the equation by the right hand side and reorganizing, we obtain 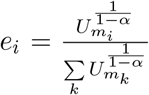. Thus, in this case, 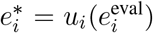 when *α →* 0. **Fig. 2d** shows that for power functions, a single evaluation can provide high performance for situations with an appreciable level diminishing marginal utility, up to about *α* = 0.5. To maintain performance in situations with weakly diminishing returns, the efficient strategy must be repeated, i.e., 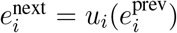 (**Fig. 2d**). This finding provides an important insight regarding diminishing marginal utility in the derivation process leading to the optimal strategy (**Fig. 4**). Specifically, IM 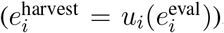, which is to approach the matching equilibrium of Eq. 2, is subjected to even stronger requirements on diminishing marginal utility than the individual fractions of the molar equilibrium. In particular, a previous study^25^ found that the same power functions considered here render Eq. 2 optimal for any 0 *< α <* 1. In comparison, the molecular IM strategy requires more strongly diminishing returns (**Fig. 2d**) to yield adequate gain.

Let us now inspect how the level of diminishing marginal utility governs the convergence of Instant Matching in a general case. Consider the iterative form of IM 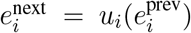, or, an equivalent formulation based on Eq. 3: 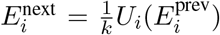. For a gain situation that exhibits strong diminishing marginal utility, 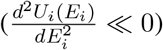, yet a gain that does not show negative returns 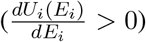, changes in *E*_*i*_ lead to only small changes in *U*_*i*_. In that case, 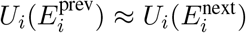 and 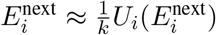. This approaches the relativistic formulation of the matching equilibrium: 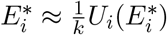 or 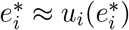. The above results generalize to situations in which each option can be characterized with a distinct function and situations in which the initial effort allocation is not equal but random (**Fig. 5b**).

### Simulations

#### Decision options

The simulations were designed to sample a large portion of the space of possible utility-effort contingencies, i.e., *U* (*E*) functions. The functions were of distinct kinds (**Fig. 5a**), used distinct magnitudes, and distinct level of diminishing marginal utility. Specifically, power, logistic, hyperbolic, and logarithmic functions in the forms 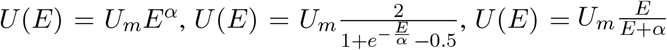, and 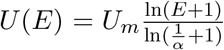, respectively, were evaluated. The logarithmic functions used *E* +1 in their argument to avoid negative utility, and ln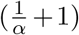 in the denominator to implement logarithm base values 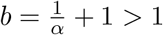.

#### Choice situations

The study simulated a large number of choice situations to test the generalizability of the findings beyond those derived analytically above. The situations featured *n* = 2, 3,… 10 options. For each option within the set of *n*, the specific *U* (*E*) function (power, logistic, hyperbolic, or logarithmic), its magnitude *U*_*m*_, and the saturation parameter *α* were all selected randomly. The function kind was selected from the set of the four. *U*_*m*_ was selected randomly from a uniform distribution over the range from 0 to 1. The diminishing marginal utility level *α* was selected randomly from the set {0.1, 0.2,. 1.0}. The randomization was performed 1000 times for each *n*, yielding a total of 9000 choice situations.

#### Stochastic choice situations

These stochastic simulations used the same number of repetitions and the same number of options. In a variable interval (VI) schedule, a binary reward is assigned to an option randomly from the Poisson process at a specific reward rate. The reward rate was again assigned randomly to each option, within the range from 0 to 1 per unit of time. A reward assigned to an option is available until the decision-maker harvests it. Harvesting events (i.e., the decision-maker’s behavior) are also binary and assigned to an option randomly from the Poisson process, now at the rate of the specific rate of effort (range from 0 to 1 per unit of time, subject to 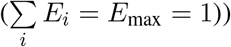. To estimate the reward rate from the noisy binary information, decision-makers must integrate the history of past rewards. A common approach for doing so is reinforcement learning, in which the integrated binary rewards are discounted exponentially^20–22^, i.e., using a function *Ae*^*-γt*^. The learning rate, is related to the time period *T* of integration. This study used 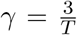, for *T* = 400. *A* was set such that the integral of the exponential over *T* was equal to one, i.e., 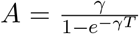. Each option evaluation operated on subsequent time period *T* ; i.e., each option evaluation encountered a distinct distribution of rewards and behaviors, generated using the respective rates.

#### Optimizing agent

In each of the above 9000 choice situations, the optimizing agent aimed to maximize its gain within a constrained distribution of effort devoted to each option. Specifically, the agent was maximizing 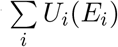 subject to 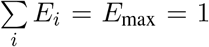. The problem was solved using constrained nonlinear multivariable optimization (function fmincon in Matlab, using the default interior-point algorithm). The algorithm converged onto a solution in a finite number of options evaluations in each decision case (see **Fig. 1** and **Fig. 3a** for the number of option evaluations necessary to reach a specific performance level).

#### Instant Matching

Like the optimizing agent, the biologically feasible decision-makers distributed their effort *E*_*i*_ across the choice options subject to finite effort, 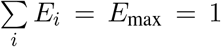. Their total gain 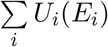, is reported as a percentage of the maximum possible gain harvested by the optimizing agent. Unlike the agent, however, the decision-makers used a single or just a small number of evaluations of the decision options. Specifically, they first sampled the worth 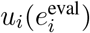 of their options *i* by allocating initial effort *e*^eval^ to each option. Subsequently, they applied the Instant Matching strategy i.e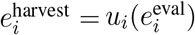. Thus, the decision-makers simply allocated their effort in proportion to an initially established worth of their options. In **Fig. 2b, Fig. 2d**, and **Fig. 5b**, this strategy was applied iteratively, i.e., 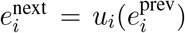, up to 4 times. Moreover, the analysis of **Fig. 2b** also tested whether the strategy is sensitive to the kind of the initial effort distribution—equal 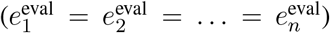 or entirely random. Given that an equal allocation of effort across options may not be realistic in natural or human settings, the results in **Fig. 2c, Fig. 2d**, and **Fig. 5b** used the random allocation of initial effort.

## Acknowledgements

I thank Drs. Alex Kacelnik, Alasdair Houston, Leo Sugrue, and Julian Brown for comments on the manuscript. This work was supported by the NIH grants R00NS100986 and RF1NS128569.

## Data and code availability

The simulation code is available and documented at neuralgate.org/download/EDM.

## Notes

### Competing Interest Statement

The authors have declared no competing interest.

### Summary of Updates

This revision reframes the paper, putting all derivations into Methods. Moreover, the paper changes the name of the strategy to Instant Matching, which is closer to the related Matching Law.

